# Somatic Cx36 gap junctions expand the anatomical substrate for electrical coupling in the adult mouse inferior olive

**DOI:** 10.64898/2026.09.09.750283

**Authors:** Kohgaku Eguchi, Mykola Medvidov, Marylka Yoe Uusisaari

## Abstract

The inferior olive (IO) is unique among adult mammalian brain nuclei in that its neurons communicate solely via gap junctions. For more than half a century, models of cerebellar timing, synchrony, and learning have therefore centered on how electrical coupling between olivary neurons is regulated. In the prevailing anatomical model, IO gap junctions reside on dendritic spines within olivary glomeruli, where nearby inhibitory terminals can locally regulate coupling through shunting conductances. Here we show that this view is incomplete. In the adult mouse IO, closely apposed neuronal somata are common, direct soma-soma appositions are visible by volume electron microscopy, Cx36 puncta occur at somatic contacts, and freeze-fracture replica labeling identifies Cx36-associated gap junction plaques on large continuous membrane surfaces. These soma-proximal gap junctions extend olivary electrical coupling beyond dendritic-spine glomeruli, challenging a core anatomical assumption in models of cerebellar coordination.

## Introduction

Among the distinctive structural and physiological features of the mammalian inferior olive (IO), inter-neuronal communication mediated exclusively by electrical synapses is particularly striking. Connexin36 (Cx36)-based gap junctions are abundantly expressed in IO neurons [1], and electrical coupling is central to prevailing theories of IO subthreshold oscillations, cerebellar complex spike synchrony, and cerebellar learning [2, 3]. Following the influential proposal by Rodolfo Llinas, the specific localization of gap junctions on dendritic spines embedded within specialized microdomains that include nearby GABAergic axon terminals has been proposed to allow rapid and reversible modulation of coupling strength by local inhibitory conductances [4]. In this framework, synaptic inhibition arising from the cerebellar nuclei can locally shunt the electrically coupled spine compartment, thereby allowing the cerebellar cortex to regulate its own activation structure through the nucleo-olivary feedback pathway [5, 6]. Direct [7, 8] and indirect [9, 10] studies have demonstrated that GABAergic synaptic activation has modulatory potency for olivocerebellar synchrony, including the emergence of olivary subthreshold oscillations proposed to gate cerebellar afferent signaling [11, 12].

Consistent with this model, ultrastructural studies have reported inferior olive gap junctions predominantly on dendritic spines, with somatic gap junctions described mainly under pathological conditions [13, 14]. Detailed 3D reconstructions have further emphasized that the spatial arrangement of IO neuronal dendrites is likely to shape how synchronized activity emerges within the nucleus [15]. As a result, electrical coupling in the adult IO is widely considered to be largely confined to spine-based glomerular microdomains. This anatomical assumption matters because the predicted effect of a gap junction depends on where it is placed. A small distal spine can be locally regulated by nearby inhibitory input, whereas a soma-proximal or thick-dendrite junction would be expected to interact with the electrotonically compact cell body and proximal dendritic tree in a different way.

Here, we re-examine the anatomical substrate of olivary electrical coupling using complementary light microscopy, volume electron microscopy, Cx36 immunohistochemistry, and freeze-fracture replica labeling (FFRL) electron microscopy. We show that adult mouse IO somata are often found closely apposed to each other with Cx36 immunolabeled puncta in between them, and that gap junction plaques reside on uninterrupted membrane areas exceeding 20 ~ *µ*m^2^, incompatible with dendritic spine morphology. These observations indicate that the adult mouse IO contains a soma-proximal electrical-coupling substrate that is not captured by the canonical spine-glomerulus model.

## Results

### Close apposition of somata is common among adult inferior olive neurons

Casual observation of IO slices in which somatic structures were revealed with Calbindin staining suggested that neuronal somata often lie extremely close to one another, with little or no intervening space (Fig. 1A). To quantify this impression, we reanalyzed a previously published dataset consisting of soma positions and perimeters obtained unilaterally from an adult mouse IO [15]. The dataset contained 8,591 somata across eight IO slices and allowed estimation of both soma diameter and three-dimensional nearest-neighbor distance (Fig. 1B).

**Figure 1:**
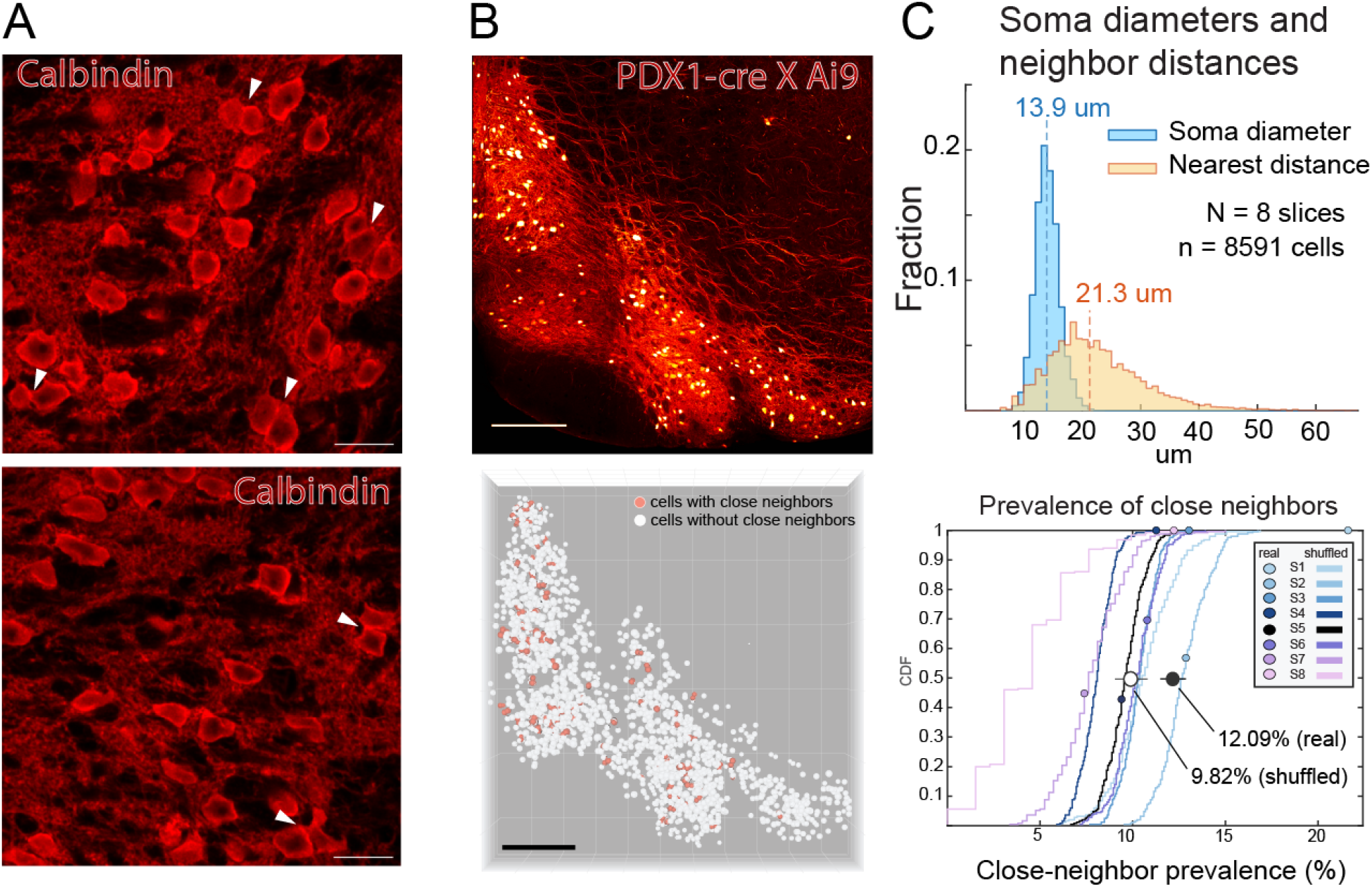
Inferior olive somata frequently have close neighbors. (A) Calbindin-labeled inferior olive somata with little intervening space (arrowheads). (B) PDX1-Cre × Ai9 dataset and soma-center map; orange points indicate somata with close neighbors. (C) Top, soma-equivalent diameters overlap the distribution of nearest-neighbor center distances, indicating that many somata lie close enough to contact. Bottom, close-neighbor prevalence was compared with packing-aware shuffled null datasets for each slice (S1–S8). Colored points show the prevalence measured from the real soma positions, and matching colored curves show the cumulative distributions obtained after shuffling soma positions within each slice footprint. The rightward displacement of real values relative to the shuffled distributions indicates that close somatic neighbors occur more often than expected from packing constraints alone. Scale bars: A, 50 *µ*m; B, 200 *µ*m.

The distribution of nearest-neighbor distances overlapped substantially with the distribution of soma diameters, indicating that many neighboring somata are close enough to be directly apposed (Fig. 1C). We classified a soma as having a close neighbor when the center-to-center distance to another soma was less than or equal to the mean of the two soma-equivalent diameters. By this criterion, approximately one in eight somata had at least one close neighbor (1,039/8,591 somata; pooled prevalence, 12.09%; per-slice range, 7.38–21.6%; Fig. 1C). To test whether this prevalence could arise from anatomical constraints alone, we generated stack-specific shuffled datasets that preserved soma diameters and *z* positions while redrawing positions within each stack footprint. The observed pooled prevalence was higher than the packing-aware shuffled null expectation (mean 9.82%; Monte Carlo *p* = 0.004), indicating that close somatic neighbors occur more frequently than expected under this spatial null model.

### Volume electron microscopy reveals direct soma-soma appositions

Light microscopy cannot resolve whether closely neighboring somata are separated by extracellular space, glial processes, or direct membrane apposition. We therefore examined the ultrastructure of IO somata using serial block-face volume electron microscopy (Fig. 2). In an approximately 100 × 100 × 12 *µ*m image stack from the principal IO, multiple somata and proximal processes could be reconstructed in near-entirety in three dimensions (Fig. 2A,B). Within this volume, we observed repeated instances of extreme close somatic apposition with no visible extracellular gap between neighboring plasma membranes (Fig. 2C).

**Figure 2:**
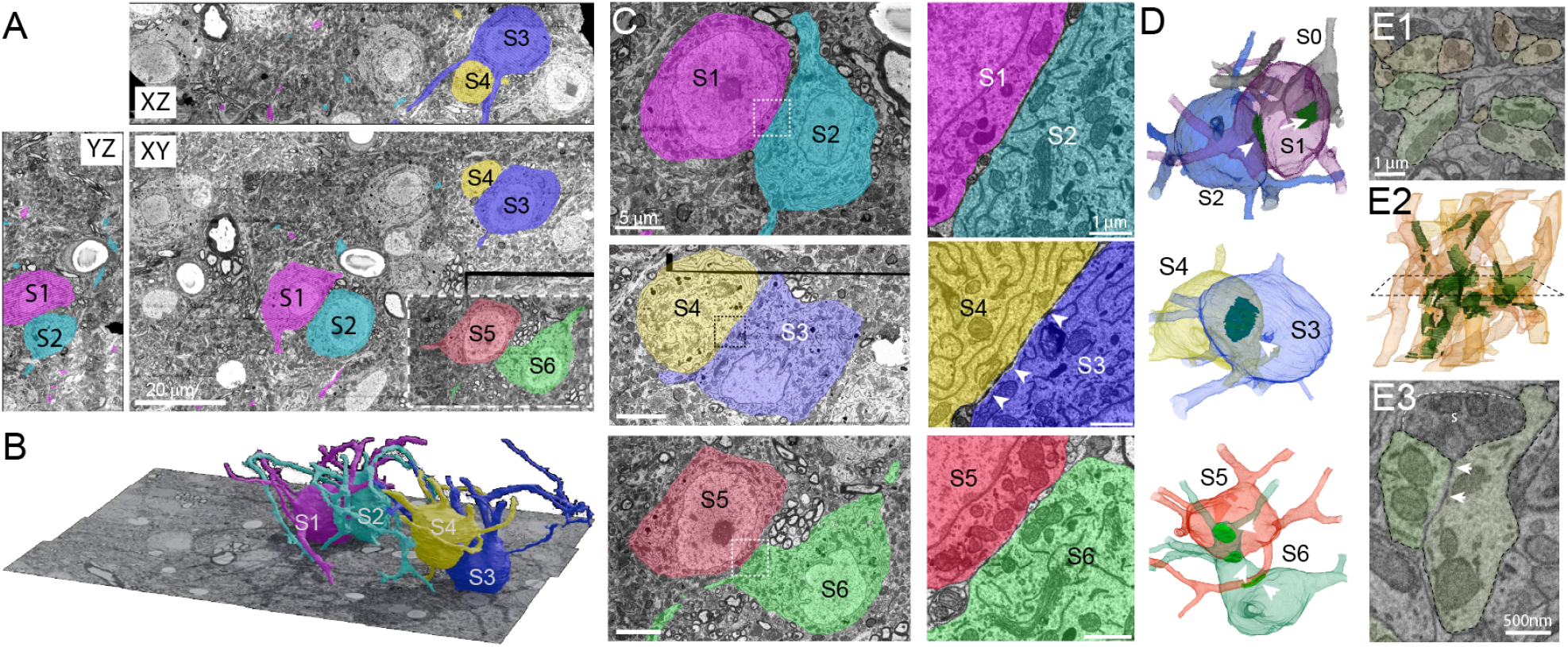
Volume electron microscopy reveals direct soma-soma appositions. (A) Orthog-onal views through a serial block-face EM volume containing olivary somata. Inset shows image plane at a different depth. (B) 3D webKnossos rendering of S1–S4. (C) Examples of closely apposed somatic membranes. Rectangular areas are shown enlarged on the right. (D) 3D visualization of reconstructed somata shown in C. Dark green plaques and white arrowheads highlight regions of somatic membrane appositions. Note that soma S0 in the top panel is incompletely contained in the volume and not shown in panels A–C. (E) Example of a dendritic “thicket” in which numerous smooth dendrites are seen in contact with each other. E1, more than 10 dendritic segments with membrane apposition with neighbors. E2, 3D visualization of manual reconstruction of the dendrites shown in E1. Dark green patches denote membrane contact areas. Dashed rectangle indicates approximate position of image shown in E1. E3, enlarged region of a three-dendrite thicket. Arrows highlight electron-dense membrane locations, possibly indicating gap junctions. Note that in the incompletely reconstructed tissue it is not possible to judge whether the dendrites belong to the same neuron. Black rectangular regions in A and C are caused by mismatches in image acquisition tiling.

The three apposed pairs of somata that were included in the volume in their entirety had membranes in contact over large plaque-like areas (Fig. 2D). Similar membrane appositions were also observed between somata and proximal dendrites (see somata S5 and S6 in Fig. 2D). The apposed somatic membranes frequently contained electron-dense puncta, reminiscent of soma-soma contact specializations reported in the suprachiasmatic nucleus [16]. Close appositions were also common between thick primary dendrites, occasionally forming dendritic “thickets” where numerous smooth dendrites shared membrane contacts (Fig. 2E1). Occasionally, electron-dense membrane regions reminiscent of gap-junctional features were observed at dendrite-dendrite appositions (Fig. 2E3). These thickets were often contacted by presynaptic terminals (example indicated “s” in Fig. 2E3), although it was not possible to reliably distinguish GABAergic from glutamatergic terminals in the volume.

Although the resolution of this volumetric EM dataset does not allow unequivocal identification of gap junction channels, the observations show that adult IO somata and proximal dendrites can provide large membrane surfaces suitable for electrical coupling, either by ephaptic effects or gap junctions.

However, membrane appositions and associated specializations do not by themselves establish the presence of gap junctions. Adherens-junction components at somatic contacts between mesencephalic trigeminal neurons persist in Cx36-null mice [17], indicating that such contacts can be maintained independently of Cx36. We therefore sought molecular and ultrastructural evidence for gap junctions at soma-proximal membranes.

### Cx36 puncta occur between closely apposed IO somata

We next asked whether molecular markers of neuronal gap junctions are present at close somatic appositions. Brainstem sections containing the IO were immunolabeled for Cx36, the predominant connexin in IO neurons [3], together with Calbindin to visualize neuronal structures. In regions where somata were closely apposed within the immunolabeled volume, up to 8 *µ*m from the slice surface, we repeatedly observed numerous Cx36 puncta located at the interface between the somata (Fig. 3). These puncta were seen against a background of additional Cx36 signal in the surrounding neuropil, consistent with the known abundance of dendritic-spine gap junctions in the IO.

**Figure 3:**
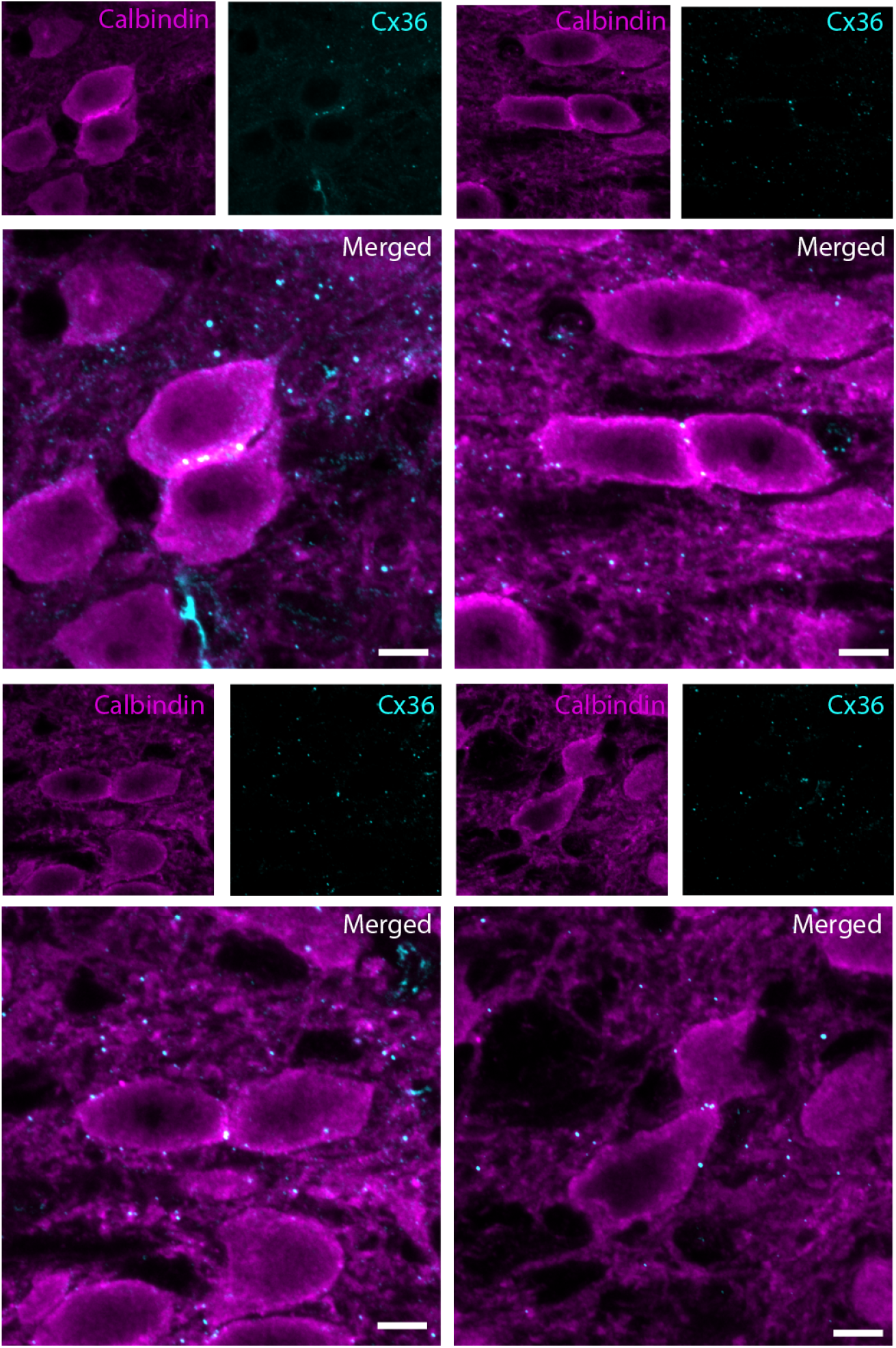
Cx36 puncta occur repeatedly at somatic appositions. Confocal images of Calbindin-labeled olivary somata and Cx36 immunolabeling from multiple examples of closely apposed somata. Images are maximum-intensity projections of optical sections acquired at 0.1 *µ*m z-steps, with each projected substack spanning less than 2 *µ*m in *z*. For each example, single-channel Calbindin and Cx36 panels show the cellular context and punctate Cx36 signal, and the merged panels show Cx36 puncta concentrated at or immediately adjacent to the soma-soma interface. The repeated occurrence of puncta at close somatic contacts supports the presence of Cx36-containing junctional specializations beyond dendritic spines. Scale bars: 5 *µ*m.

The confocal observations do not by themselves prove that the puncta are somatic gap junction plaques, because light microscopy cannot resolve the membrane topology of small structures near soma surfaces. In the absence of controlled observations that could provide reliable estimation of labeling efficiency and specificity, we also did not attempt to quantify gap junction prevalence from these sections. However, the observations bridge the light-microscopy evidence for close somatic packing with the electron-microscopy evidence for direct soma-soma contact, and support the presence of Cx36-containing junctional specializations beyond dendritic spines.

### Freeze-fracture replica labeling identifies gap junction plaques on large membrane structures

To obtain direct ultrastructural evidence for gap junctions on soma-sized membrane surfaces, we acquired freeze-fracture replica labeling images from adult mouse IO (Fig. 4A). As expected, we observed gap junction plaques on thin neurites, often near synaptic structures (Fig. 4B). However, we also found gap junction plaques scattered across large continuous membrane areas exceeding 20 *µ*m^2^ (Fig. 4C). These uninterrupted membrane areas were too large to be compatible with dendritic spine morphology and were consistent with somatic or proximal dendritic membranes.

**Figure 4:**
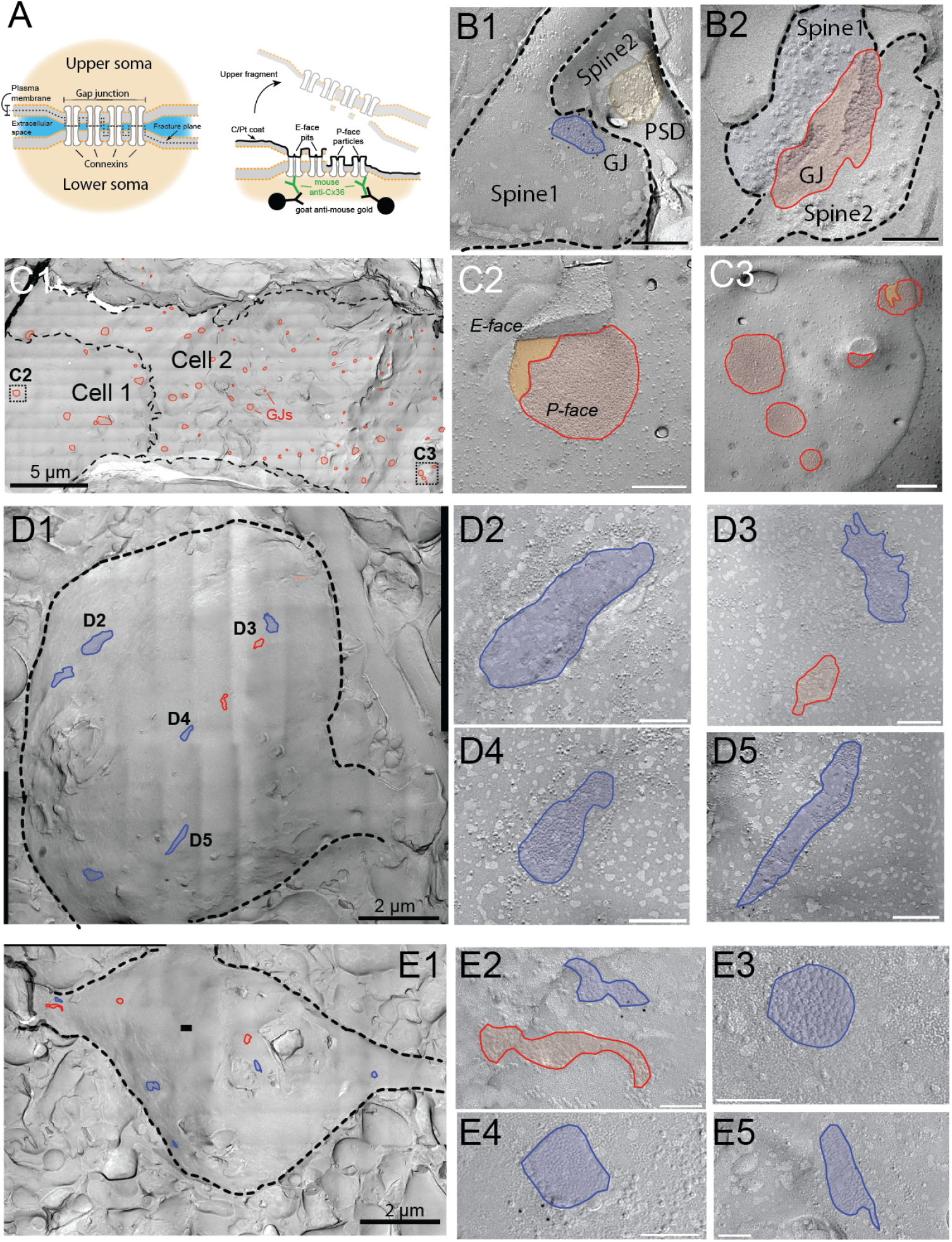
Freeze-fracture replica labeling identifies somatic Cx36 gap junctions. (A) Schematic of freeze-fracture replica immunogold labeling of gap junctional plaques. (B) Gap junction plaques on spine-like processes. Immunogold labeling in the left panel demonstrates Cx36. Scale bars: 200 nm (left) and 100 nm (right). (C) Gap junction plaques (red) on large continuous membrane areas consistent with somata. Inset locations are enlarged in C2–C3. E- and P-faces are indicated. Inset scale bars: 200 nm. (D,E) Cx36 immunogold-labeled gap junction plaques (blue) on large membrane areas. Insets (D2–D5, E2–E5) are enlarged. Plaques without visible immunogold labeling are shown in red. Inset scale bars: D, 200 nm; E, 200 nm.

In the IO, gap junctions are known to be present both on neuronal and glial membranes [18, 19]. To rule out the possibility that the large membrane structures with identified gap junctions represented glial cells rather than IO neurons, we conducted additional FFRL experiments with immunogold labeling for Cx36, which is known to be present on neurons [20]. In these samples, we recovered numerous examples of putative somatic membranes carrying gap junction plaque structures associated with gold particles (Fig. 4D,E). Further support for the neuronal identity of the membrane came from the round and regular particle arrangement of the plaques, distinguishing them from the characteristic elongated glial plaques formed by glial connexins [20].

The putative somatic gap junction plaques were on average around three times larger than gap junctions found on spines (spine: 0.023 ± 0.002 *µ*m^2^, n = 29 GJs; soma: 0.070 ± 0.006 µm^2^, n = 75 GJs from 11 somata; mean ± s.e.m.). Notably, they were in some cases seen aligned in “bands” across the somatic membrane, suggesting a possible somatic contact with a proximal dendrite passing by, as seen in the example shown in Fig. 2D (S5–S6). These observations establish that adult mouse IO gap junctions are not restricted to dendritic spines.

## Discussion

Immunogold particles near plaques on large continuous membranes provided direct evidence for Cx36-associated gap junctions on IO somata or soma-sized membrane surfaces. These observations establish that adult mouse IO gap junctions are not restricted to dendritic spines and revise the prevailing view of electrical coupling in the adult mouse inferior olive. Soma-proximal olivary gap junctions are not entirely without precedent: De Zeeuw and colleagues reported gap junctions between a soma and a primary dendrite in the dorsal cap of Kooy [21]. However, the prevailing adult IO model has remained dominated by spine-glomerulus coupling, where nearby inhibitory synapses can regulate coupling strength by local shunting.

While we do not currently have the means to examine the functional contribution of somatic gap junctions to olivary physiology, these anatomical observations have implications by themselves. Unless the connexin channels are predominantly closed, large gap junction structures residing on voluminous neuronal compartments such as somata are likely to create significant synchronicity of sub- and suprathreshold electrical behavior. Strikingly, it could be presumed that the output of neurons coupled at their somata is to lose independence, especially if the axon emerges close to the site of coupling. In this context it is curious to consider that the axon commonly stems from a dendrite in the IO [14].

Another implication for olivocerebellar dynamics regards the question of modulation of inter-neuronal signaling within the olive. In contrast to gap junctions embedded in distal spine glomeruli, where GABAergic synaptic conductances originating from the cerebellar feedback pathway can generate sufficient conductance to significantly decrease effective electrical coupling, it is not clear to what extent a similar process could modulate somatic or thick-dendrite coupling. Notably, while we observed large presynaptic terminals close to dendro-dendritic contact points (Fig. 2E3), nucleo-olivary terminals only sparsely target IO somata [7]. Hence, a distinct mechanism for regulating somatic coupling needs to be identified. One possible mechanism involves glutamatergic regulation of olivary coupling [22, 23], as NMDA receptor subunit immunoreactivity has been reported in somata and proximal dendrites of IO neurons [24].

Indeed, even though gap junctional coupling can be significantly suppressed by GABAergic synaptic conductances, the coupling is not completely abolished [7], and residual coupling may be mediated by soma-proximal gap junctions.

Finally, we must raise the question regarding the lack of earlier observations. If the somatic gap junctions indeed are common, why were they missed in the past? We believe the answer lies in the technical limitations of the classical approaches. In conventional thin-section electron microscopy, large membrane appositions are difficult to recognize without volumetric context, and a gap junction plaque on an extended membrane surface can easily be missed if the sectioning plane does not intersect it. By contrast, freeze-fracture replica labeling reveals intramembrane particle arrays directly across broad membrane surfaces. Immunohistochemical evidence is likewise constrained by sampling geometry. Even if approximately one in eight IO somata has a close neighbor in situ, and even if every such apposition contains a gap junction, only the small subset of contacts located within the efficiently antibody-penetrated surface region of the slice would be detectable. Consistent with this constraint, even in sections examined specifically for soma-soma contacts, we typically found only a few suitable appositions per slice.

Physiological detection is also expected to be difficult. Although somatic gap junctions might be expected to produce substantially larger coupling ratios than those reported in numerous electrophysiological studies [7, 25, 26], paired recordings from coupled neighboring IO somata would be extremely rare in acute slices unless such pairs were specifically sought. It is also possible that under experimental conditions only a small fraction of somatic connexin channels are open. Consistent with this possibility, somatic Cx36-based gap junctions in mesencephalic trigeminal ganglion neurons showed very low compound junctional conductance *in vitro*, implying that most channels were closed [27].

In all, we believe that the present findings capture a genuine anatomical feature of the adult mouse inferior olive rather than a technical or sampling artifact. Granted, the gap junctions on dendritic spines remain an elegant substrate for rapid modulation of coupling by local synaptic conductances. However, the anatomical evidence presented here calls for broadening the view on olivary network communication to include soma-proximal contacts. Our spatial analysis estimates that approximately one in eight IO somata has a close somatic neighbor, although this does not establish the prevalence of somatic gap junctions. Appositions involving proximal dendrites are likely much more common and may provide substantial membrane contact for electrical interactions, potentially including ephaptic signaling. This possibility may be relevant when considering why some coordination of olivary activity appears to persist even in Cx36 knockout mice [3, 28, 29].

Going further, we would like to reverse the question: are the weak coupling coefficients observed in vitro [7, 26, 30] truly aligned with the high levels of olivary spike coactivation observed in living animals [8, 31]? To the point, large-scale computational models reproducing olivary dynamics tend to abstract the dendritic structures [32–34]. Hence, the presence of somatic gap junctions may be more consistent with the implied assumptions of these models than with coupling restricted to electrically remote spines.

Finally, while somatic gap junctions may be a special adaptation to the unique physiology of olivary neurons, the fact that they have been overlooked across decades of work raises the possibility that similar soma-proximal electrical synapses may be more common in adult mammalian brain than currently appreciated.

### Limitations of the Study

The present study does not directly measure electrical coupling through somatic gap junctions. The conclusion is anatomical: Cx36-associated gap junction plaques occur on soma-sized or proximal large membrane structures in the adult mouse IO. Functional experiments targeted to anatomically identified soma-soma or soma-dendrite pairs will be required to determine when these junctions are open, how strongly they couple the cells, and whether they are regulated by the same or different mechanisms as spine-based junctions.

## Methods

### Animals

Male C57BL/6J mice aged 2–5 months were used in accordance with the 2006 guidelines for Proper Conduct of Animal Experiments of the Science Council of Japan and protocols approved by the Okinawa Institute of Science and Technology Animal Care and Use Committee (#2024-009). Data used for the soma-neighbor analysis in Fig. 1 were from a previously completed study [15].

### Soma-neighbor analysis

Curated FIJI ROI tables from eight IO image stacks were analyzed. Each soma was represented by centroid coordinates (*x, y, z*), with *z* computed from slice number using a 2 *µ*m interval. Soma diameter was estimated as the equivalent-circle diameter from traced area. For each soma, 3D center-to-center distances to all other somata in the same stack were computed. A soma was classified as having a close neighbor if another soma lay within the mean of the two soma diameters. To estimate the prevalence expected from packing constraints alone, a packing-aware spatial null distribution was generated by preserving soma diameters and *z* positions while redrawing *x, y* positions within the convex hull of the observed soma centers in each stack. Candidate positions were sampled uniformly within this footprint and rejected if they placed a soma center at a three-dimensional distance from an already placed soma that was smaller than the larger of the two soma radii. Null prevalence was estimated from 250 shuffled datasets per stack using random seed 1. Pooled prevalence was calculated from the total number of somata with close neighbors across all stacks divided by the total number of somata. The one-sided Monte Carlo *p*-value was calculated as (1 + *k*)*/*(1 + *N*), where *k* is the number of pooled shuffled prevalences greater than or equal to the observed pooled prevalence and *N* = 250.

### Confocal immunohistochemistry

Mice were anesthetized with phenobarbital by intraperitoneal injection (400 mg/kg) and transcardially perfused with PBS for 1 min, followed by 4% paraformaldehyde (PFA) in PBS for 5 min at room temperature. Brains were removed, cryoprotected in 30% sucrose in PBS overnight, embedded in tissue-tek O.C.T. Compound, frozen at −80°C, and sectioned coronally at 50 *µ*m with a cryostat (CM1950, Leica). Unless otherwise noted, washes were performed for 10 min at 80 rpm. Brainstem sections containing the IO were washed three times in PBS and blocked for 1 h at room temperature in PBS containing 10% bovine serum albumin (BSA), 1% normal goat serum, and 0.4% Triton X-100. Sections were incubated at 4°C for 48 h in antibody buffer (2% BSA and 0.4% Triton X-100 in PBS) containing mouse anti-Cx36 primary antibody (1:200 dilution, 2.5 *µ*g/mL, #37-4600, Thermo Fisher Invitrogen) and guinea pig anti-calbindin antibody (1 *µ*g/mL, #214 005, Synaptic Systems), with shaking at 80 rpm. After three washes in PBS, sections were incubated in the same buffer containing Alexa Fluor 488-conjugated goat anti-mouse IgG and Alexa Fluor 594-conjugated goat anti-guinea pig IgG secondary antibodies (1:1000 dilution, A-11029 and A-11076, Thermo Fisher Invitrogen) at 4°C for 17 h with shaking at 80 rpm. Sections were washed, mounted on glass slides, and coverslipped with Vectashield antifade mounting medium (H-1900, Vector Laboratories). Images were acquired with a Zeiss LSM 880 laser-scanning confocal microscope in line-scan mode with 16-bit depth. Cx36 was excited with a 488 nm argon laser and emission was collected between 493 and 584 nm. Calbindin was excited with a 594 nm HeNe laser and emission was collected between 599 and 734 nm. Overview tiled images and z-stacks were acquired with a 20× objective (Plan-Apochromat 20×/0.8 M27) and stitched in ZEN (Zeiss). High-magnification z-stacks were acquired with a 63× objective (Plan-Apochromat 63×/1.4 Oil DIC M27) at 0.1 *µ*m z-steps. Maximum-intensity projections were generated in Fiji.

### Volume electron microscopy

A C57BL/6J male mouse was anesthetized by intraperitoneal injection of MMB anesthesia (0.1 mL/10 g body weight; medetomidine hydrochloride 1 mg/mL, 0.75 mL; midazolam 10 mg/2 mL, 2 mL; butorphanol tartrate 5 mg/mL, 2.5 mL; saline, 19.75 mL) and transcardially perfused with PBS followed by 2.5% glutaraldehyde (Sigma, G7776-10ML) and 2% formaldehyde (Thermo Fisher, 28908) in 0.1 M phosphate buffer (pH 7.4). Brains were dissected and postfixed overnight in the same fixative. Sections were cut rostrocaudally at 200 *µ*m with a Leica VT1000 S vibratome and contrasted for serial block-face imaging using an OTO staining protocol [35]. Sections were contrasted with 2% osmium tetroxide and 3% potassium ferrocyanide in 0.15 M sodium cacodylate buffer for 1 h at room temperature, rinsed in water, incubated with 1% thiocarbohydrazide for 20 min, rinsed, contrasted with 2% osmium tetroxide for 30 min, rinsed, incubated in 1% uranyl acetate overnight at 4°C, rinsed, and contrasted in Walton’s lead aspartate solution for 30 min at 60°C.

Samples were dehydrated through ethanol and acetone, infiltrated with graded Epon resin, embedded in flat molds, and polymerized for 48 h at 60°C. Sections were mounted on the microscope sample holder with superglue, polished with an Apreo VS diamond knife (Diatome), and imaged with an FEI Teneo VolumeScope scanning electron microscope. An area containing the principal IO was acquired as a 3D dataset with 60 nm section thickness, 8 nm pixel size, 500 ns pixel dwell time, 200 nA probe current, 2 kV accelerating voltage, 1,378 sections, and a 6 × 4 grid of 8192 × 8192 pixel tiles with 10% overlap. Image tiles were aligned, and neuronal somata, proximal processes, and apposed membrane regions were manually reconstructed in webKnossos.

### Freeze-fracture replica labeling

Mice were anesthetized with phenobarbital by intraperitoneal injection (400 mg/kg) and transcardially perfused with PBS for 1 min followed by freshly prepared 2% PFA in 0.1 M sodium phosphate buffer (pH 7.4) for 10 min at room temperature. Coronal brainstem slices (150 *µ*m thick) containing the IO were cut in PBS with a vibratome (5100mz, Campden Instruments) and cryoprotected sequentially in 10%, 20%, and 30% glycerol in 0.1 M sodium phosphate buffer. A total of nine mice were processed for freeze-fracture replica preparation. IO-containing tissue blocks were dissected with a microscalpel (#10316-14, FST) and frozen with a high-pressure freezer (HPM100, Leica). Frozen samples were fractured at −100 to −120°C and replicated by sequential deposition of carbon (3–5 nm), carbon-platinum (2 nm, unidirectionally from 60°), and carbon (20–25 nm) in a freeze-fracture machine (JFD-V, JEOL). Samples were digested in 2.5% SDS in 0.1 M Tris-HCl (pH 8.3) at 70°C for 20–22 h.

Unless otherwise noted, replica washes were performed for 10 min. Replicas were washed once in SDS solution and twice in washing buffer containing 0.1% fatty acid-free BSA in 50 mM Tris-buffered saline (TBS, pH 7.4). For Cx36 immunolabeling, replicas were blocked for 1 h at room temperature in TBS containing 3% BSA, 2% cold fish skin gelatin, and 0.05% Tween-20, then incubated with mouse anti-Cx36 primary antibody (1:100 dilution, 5 *µ*g/mL, #37-4600, Thermo Fisher Invitrogen) diluted in 1% BSA, 1% cold fish skin gelatin, and 0.05% Tween-20 in TBS at 15°C for 1–2 overnights. After three washes in washing buffer, replicas were incubated with 6 nm colloidal gold-conjugated goat anti-mouse IgG secondary antibody (1:20 dilution, #115-195-146, Jackson ImmunoResearch) in the same buffer at 15°C for 17 h. Replicas were washed twice in washing buffer and then in distilled water, mounted on formvar-coated grids, and imaged with a JEOL JEM-1400 Flash transmission electron microscope operated at 100 kV at 30,000× to 50,000× magnification. Gap junctions were identified from Cx36 immunolabeling together with established ultrastructural criteria, including regular intramembrane particle arrays on P-faces, corresponding pits on E-faces, and narrowing of the extracellular space at junctional borders [36].

### Image analysis

Confocal microscopy images were processed in FIJI [37]. For Fig. 3, substacks around Cx36 puncta were selected manually, channels were separated, and histograms were adjusted to visualize the signals without saturation. Three-dimensional Gaussian blurring was applied with settings of 1, 1, and 2 pixels in the *x, y*, and *z* directions, respectively, before channel merging and maximum-intensity projection. Volume-EM image stacks were aligned, stitched, manually reconstructed, and visualized using webKnossos [38]. Three-dimensional reconstruction images in Fig. 2D were rendered with MeshLab [39]. FFRL panoramic images were acquired using the limitless panorama function of the JEM-1400 Flash. Panoramic images were stitched and cropped using custom in-house napari plugins. Plaque regions of interest were manually traced, and plaque areas were measured from calibrated images. Putative spine and putative somatic plaques were classified by the membrane context visible in the replica: small spine-like processes in proximity to synaptic structures were classified as spine-associated, whereas plaques on large uninterrupted membrane areas were classified as soma- or proximal-dendrite-associated.

## Data and Code Availability

Data repositories are being prepared and are not yet publicly available. Image data will be deposited in the BioImage Archive (light microscopy and FFRL) and EMPIAR (volume electron microscopy). Raw data tables supporting the analyses will be deposited in Zenodo. Custom analysis code will be made available through GitHub. Repository links and accession numbers will be provided upon publication of the final journal article.

## Acknowledgments

We thank the OIST Animal Resources Section (ARS) for animal care and the OIST Imaging Section for microscopy and electron microscopy support.

## Author Contributions

K.E. and M.Y.U. designed research; K.E. performed experiments; K.E. and M.Y.U. analyzed data;

M.M. acquired volume electron microscopy images; and K.E. and M.Y.U. wrote the paper.

